# Invasion history of *Aedes* (Stegomyia) *albopictus* into Mesoamerica based on mitogenomes and *Wolbachia* symbionts: Multiple introductions with temperate origins

**DOI:** 10.64898/2026.07.08.737237

**Authors:** K. L. Bennett, Thomas L. Schmidt, Jonathan P. Day, José Manuel Gutiérrez Alvarado, Gabriela Delgado, Rodrigo Marín Rodríguez, Luis Fernando Chaves, Julie I. R. Labau, Owen W. McMillan, Francis Jiggins, Jose R. Loaiza

## Abstract

The global invasion of the Asian tiger mosquito *Aedes albopictus* has led to an increase in arboviral disease, including within Mesoamerica. Understanding vector invasion routes is important for public health because it directs biosecurity and identifies sources of adaptive allele spread. Panama is an important hub of global trade with opportunities for *Aedes* introduction through both maritime and overland routes but dispersal into the Isthmus has not yet been investigated. We therefore sought to investigate the population structure and invasion history of *Ae. albopictus* into Panama, targeting both its mitogenome and associated *Wolbachia*. Historical demographic analysis with Bayesian phylogeographic diffusion models and estimates of divergence revealed that Panamanian *Ae. albopictus* and its associated *Wolbachia* have a convergent evolutionary history resulting from multiple introductions. Both could be traced to Asian-derived lineages introduced via the Americas, with invasion primarily through the maritime trade of the Panama Canal rather than overland dispersal from neighboring Costa Rica. An investigation of the relative density of *Wolbachia* in Panama revealed that both the strains *w*AlbB and *w*AlbA were at a notably lower density compared to other worldwide locations. This finding has implications for arbovirus transmission and raises important questions about how *Wolbachia* density is impacted by the environment and impacts on population control. Overall, the Panama Canal is a key route for vector introductions into Mesoamerica.

## Introduction

Global dispersal of invasive mosquito vectors has intensified in recent decades, driven by human-mediated transport, climate change, and urbanisation. A prime example is the Asian tiger mosquito, *Aedes albopictus*, which has rapidly expanded from Southeast Asia since the 1980s to colonize the Americas, Europe, Africa, and the Middle East(Kraemer et al., 2015; Swan et al., 2022). This invasion is fueled by the species*’* wide ecological niche, desiccation resistant eggs (Kraemer et al., 2015), and the global trade of man-made containers such as used tyres (Swan et al., 2022). This geographic expansion has triggered a resurgence of dengue (Kobayashi et al., 2018; Luo et al., 2017; Rezza, 2012; Sacco et al., 2024) and chikungunya viruses (Faye et al., 2024; Gossner et al., 2018; Miller & Loaiza, 2015). Once regarded as a secondary vector, *Ae. albopictus* now drives chikungunya transmission owing to a viral mutation that enhances replication within the host (Tsetsarkin et al., 2007), highlighting a global need to better understand its underlying dispersal pathways.

Mesoamerica has seen a recent upsurge in these arboviral diseases (Robert et al., 2020; Segura et al., 2021). However, regional dispersal routes for *Ae. albopictus* remain poorly characterized due to sparse surveillance records (Swan et al., 2022) and limited genetic data(Battaglia et al., 2022; Eskildsen et al., 2018; Pech-May et al., 2016). As the southern-most country in Mesoamerica, and a primary global trade hub, Panama represents a key juncture for *Aedes* introduction via the Panama Canal. Although the spread of invasive *Ae. albopictus* across Panama is well documented (Bennett et al., 2019; Bennett et al., 2021a; Eskildsen et al., 2018), the exact sources and number of introductions remain unclear. Initial entomological records from Juan Diaz in ∼2002, on the outskirts of Panama City, suggest an introduction via the Panama Canal. Maritime invasion could have occurred through the commercial corridor linking the United States to East Asian manufacturing hubs in China, Japan and South Korea. Shipping includes the global trade of new and used vehicle tires, a commodity well-documented to facilitate *Aedes* transport, acting as artificial conduits for high-volume gene flow that bypasses natural geographic barriers (Bennett et al., 2019; Swan et al., 2022). Alternatively, a separate overland introduction from neighboring Costa Rica, which is the only country connected to Panama by highways has also been suggested. This is supported by later records of *Ae. albopictus* in Western Panama while remaining absent from the Azuero Peninsula in Central Panama (Miller & Loaiza, 2015).

The mitochondrial genome (mitogenome) is widely used to resolve phylogeography due to its haploid nature, high mutation rate and maternal inheritance, which collectively allow the resolution of recently diverged lineages. In *Ae. albopictus*, cytochrome oxidase I (*COI*) sequence data have revealed two distinct mitochondrial haplogroups in Panama, consistent with multiple introductions via maritime and overland routes(Eskildsen et al., 2018). Nevertheless, the robustness of these inferences is constrained by small sample sizes from limited geographic locations and restricted sequence length. Complete mitogenomes offer vastly enhanced resolution and have been successfully used to reconstruct the invasion history of *Ae. albopictus* in Asia (Battaglia et al., 2022; Schmidt et al., 2026). *Ae. albopictus* is naturally infected with two strains of the maternally inherited endosymbiotic bacterium *Wolbachia, w*AlbA and *w*AlbB. As these endosymbionts are coinherited alongside the mitochondria, they are expected to share a congruent evolutionary history with the host mitochondrial lineage since the establishment of the co-infection (Schmidt et al., 2026). Although natural selection can occasionally distort these genetic signals (Hurst & Jiggins, 2005), global baseline data demonstrate that tracking these parallel maternal markers largely reflects the historical migration pathways of the mosquito host. Characterizing both genomes simultaneously therefore provides an independent, high-resolution validation system to map the introduction routes of *Ae. albopictus* into Panama. *Wolbachia* provides a fitness advantage to infected female mosquitoes via the process of cytoplasmic incompatibility, where crosses between *Wolbachia*-infected males and uninfected females produces embryonic lethality. *Wolbachia* can also reduce the ability of mosquitoes to transmit arboviruses. This effect is greater when *w*AlbB is transferred to *Ae. aegypti* than in the natural host (Ritchie et al., 2018; Zheng, Zhang, Li, Yang, Wu, Liang, Liang, Pan, Hu, Sun, et al., 2019). This has led to control programs releasing *Wolbachia-*infected *Ae. aegypti* into wild populations to block dengue transmission. Since the efficacy of these interventions depends on *Wolbachia-* induced fitness effects and the density of *Wolbachia* in mosquito tissues, an understanding of these traits for naturally occurring strains is valuable for successful implementation(Zheng, Zhang, Li, Yang, Wu, Liang, Liang, Pan, Hu, Sun, et al., 2019).

Given the role of Panama as an important gateway for biological invasions of disease vectors into Mesoamerica, identifying the geographic origins of *Ae. albopictus* alongside the characteristics of its *Wolbachia* symbionts is vital for designing targeted vector control interventions. We therefore sought to investigate the population structure and invasion history of *Ae. albopictus* into Panama, targeting both its mitogenome and associated *Wolbachia*. We hypothesise that *Ae. albopictus* has invaded Mesoamerica multiple times, entering via maritime shipping routes through the Panama Canal from North American and Asian sources, or via overland dispersal along major international highways. We hypothesize that the genetic signals of the maternally inherited *w*AlbA and *w*AlbB *Wolbachia* strains in Panama will show evolutionary congruence with the *Ae. albopictus* host mitogenomes. Consequently, the *Wolbachia* genomic data will reflect the same parallel maternal marker trends seen in the host, providing an independent, high-resolution validation of multiple independent introduction events into the region.

## Methods

### Mosquito Sampling and Study Sites

Immature stages of *Ae. albopictus* (i.e., eggs, larvae, and pupae) were sampled across Panama using oviposition traps (ovitraps) in 2016 and 2017. To ensure sampling independence, three ovitraps were deployed at least 300 m apart at each collection location, following previously described protocols (Bennett et al., 2021). Field collections were conducted during the rainy season (May–November) of 2017 across 12 distinct locations spanning six provinces (Figures 1 and 3). The sampling design targeted high-risk entry points, including areas within Panama Province adjacent to the maritime shipping ports of the Panama Canal, as well as regions bordering Costa Rica (Chiriquí Province) and Colombia (Darién Province).

**Figure 1.**
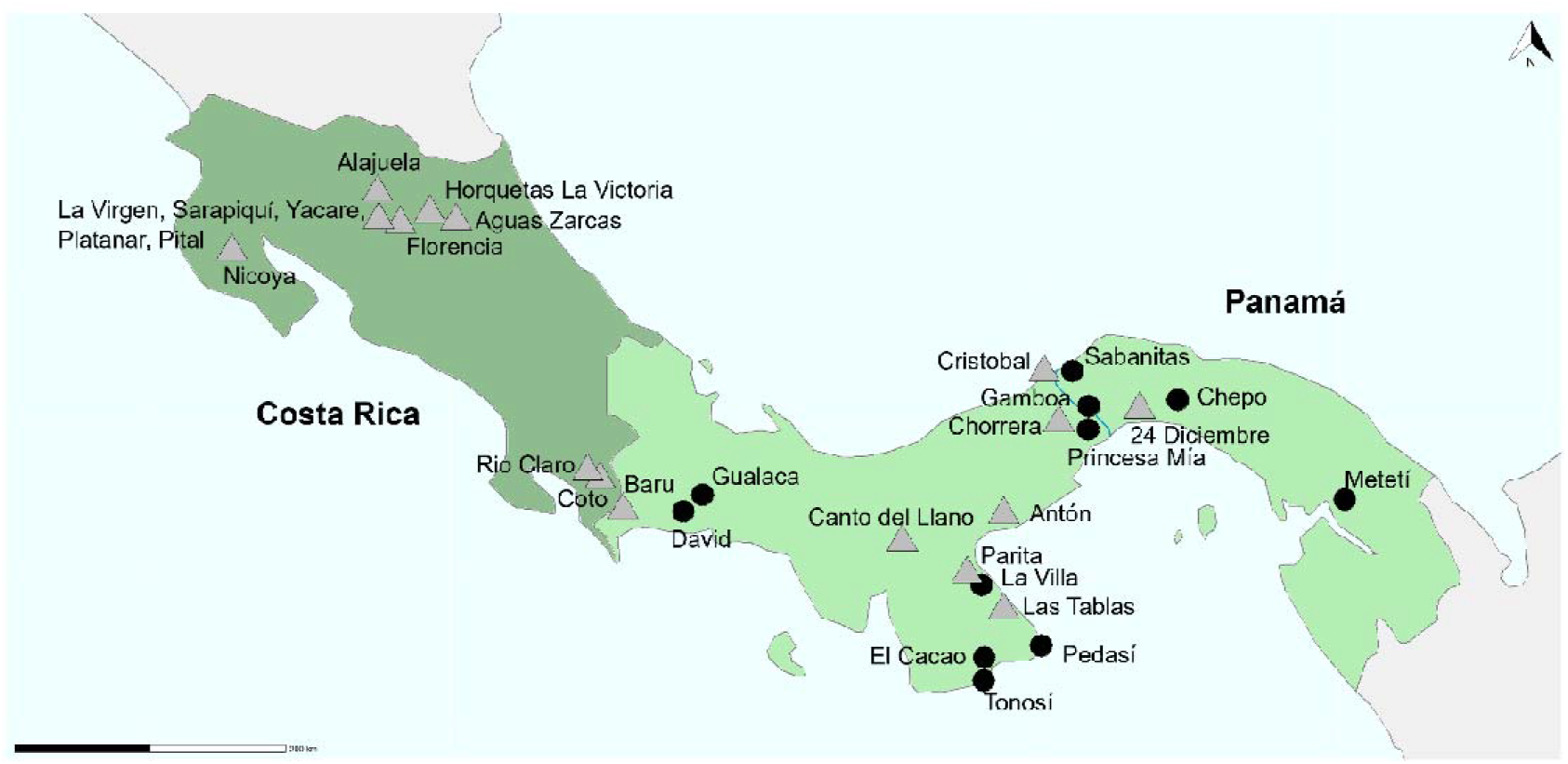
Sampling locations of *Ae. albopictus* across Panama and Costa Rica targeted either for the generation of both mitogenomes and *COI* mitochondrial sequences (black circles) or *COI* Sanger sequences only (grey triangles).

Collected specimens were transported to the laboratory, where larvae were reared to adulthood in separate containers corresponding to each ovitrap. Adult mosquitoes were morphologically identified as *Ae. albopictus* using standard taxonomic keys (Rueda, 2004) and preserved in absolute ethanol at -20 °C until molecular analysis. To complement the Panamanian dataset and contextualize regional and historical vector movement, *Ae. albopictus* was additionally collected across Costa Rica between 2019 and 2020. Larvae were collected via active surveys of natural and artificial breeding containers across 16 geographically widespread locations. These specimens were similarly preserved in absolute ethanol, transported to the processing laboratory at INDICASAT AIP in Panama City, and stored at -20 °C until analysis.

### High-Throughput Library Preparation and Target Enrichment

To avoid the sampling of siblings, one individual mosquito from each of 68 oviposition traps set up in Panama was selected for sequencing. Library preparation and target enrichment was performed as previously detailed(Schmidt et al., 2026). In brief, DNA sequencing libraries were prepared using the NEBNext® Ultra™ II FS DNA Library Prep Kit for Illumina (New England Biolabs, E7805L) and NEBNext® Multiplex Oligos for Illumina® (New England Biolabs, E7335G), yielding at least 1µg of DNA in the library for target capture. Modifications to the protocol are detailed in (Schmidt et al., 2026). Libraries were quantified using a Qubit HS DNA assay kit (Thermofisher Q32851) and the concentration assessed using a QuantStudio™ 5 Real-Time PCR System (Applied Biosystems A28575) in replicate. Equimolar quantities of libraries were combined for enrichment in pools representing 22-24 *Ae. albopictus* mosquitoes. Pooled samples were enriched for *Wolbachia* and mitochondria target DNA using a KAPA Hypercapture kit (Roche 09075810001) and KAPA Hypercapture Bead kit (Roche 09075780001) with custom designed KAPA Hyperexplore Max probes (Roche 09063633001). Target captures were quantified with qPCR and combined before sequencing on the Illumina NovaSeq platform.

### Bioinformatic Alignment and Variant Genotyping

Sequences were concurrently aligned to five genomes: *Ae. albopictus* mtDNA (Zhang et al., 2016), *Wolbachia* strains *w*AlbA (Martinez et al., 2022) and *w*AlbB (Sinha et al., 2019), and plasmids pWALBA1 and pWALBA2 (Martinez et al., 2022). Alignment was with bowtie2 (Langmead & Salzberg, 2012) using very-sensitive settings. Following alignment, we used samtools v1.7 (Li et al., 2009) to generate sorted bam files, then Picard v2.27.4 (http://broadinstitute.github.io/picard/ ) to add read groups and mark duplicates, then samtools to retain only reads mapped as primary alignments in proper pairs.

Genotyping of samples was with GATK v4.2.6.1 (Van der Auwera & O’Connor, 2020). HaplotypeCaller was first used to produce gVCF files for every sample, setting EMIT_ALL_CONFIDENT_SITES. GenomicsDBImport was used to build these into a database. GenotypeGVCFs was used to genotype all samples concurrently, with settings to “--add-output-vcf-command-line”, “--include-non-variant-sites”, and “-- sample-ploidy 1”. Hard filtering was applied to variants as follows: SelectVariants to remover indels; VariantFiltration for standard GATK hard filters (QD < 2.0, SOR > 3.0, FS > 60.0, MQ < 40.0, MQRankSum < -12.5, ReadPosRankSum < -8.0); and SelectVariants to “--set-filtered-gt-to-nocall TRUE”.

### DNA Extraction and *COI* Sanger sequencing

Genomic DNA was extracted from individual whole larvae (n = 33) from Costa Rica using the DNeasy Blood & Tissue Kit (Qiagen, Hilden, Germany) following the manufacturer’s protocols. Polymerase chain reaction (PCR) amplification targeting the mitochondrial cytochrome c oxidase subunit I (COI) gene was performed following the same procedures as previously described (Eskildsen et al., 2018). Resulting ∼658-bp PCR amplicons were purified using ExoSAP-IT (USB Corporation, Cleveland, USA) (Bell, 2008) and submitted for bidirectional Sanger sequencing at Macrogen Inc. (Seoul, South Korea). Forward and reverse chromatograms were inspected, trimmed, and assembled into consensus sequences. Multiple sequence alignments were executed using the MUSCLE and ClustalW algorithms implemented in Geneious v.7.1 (http://www.geneious.com/). All generated sequences were deposited in GenBank under accession numbers XXXXXX–XXXXXX.

### Population structure and Phylogenetic Analysis

To compare our findings to the previously described population structure of the region, the *COI* partition was extracted directly from our newly generated complete mitogenomes. This was achieved by aligning the complete mitogenomes against reference *Ae. albopictus* COI sequences from Panama (Eskildsen et al., 2018) using the MUSCLE algorithm implemented in Jalview v.2 (Edgar, 2004; Waterhouse et al., 2009). We then constructed a Neighbor-Joining phylogenetic tree using MEGA v.11 (Tamura et al., 2021) to analyze the newly extracted mitogenome sequences alongside published Panamanian sequences mined from GenBank and our newly generated Sanger sequences from Costa Rica, replicating the phylogenetic approach of Eskildsen et al. (2018). The resulting trees were functionally visualized and annotated using iTOL (Letunic & Bork, 2021).

### Historical Demography and Coalescent Divergence Estimation

A genealogical tree was generated in Beast X v10.5.0 (Drummond et al., 2012; Suchard et al., 2018) using a HKY substitution model, a constant size coalescent and a uniform clock. Independent substitution models were set for 1^st^ codon positions, 2^nd^ codon positions, 3^rd^ codon positions, non-coding RNA, and non-coding intergenic DNA partitioned using the strand (+/-) to identify the start and end of each coding sequence. Following (Schmidt et al., 2026)we generated a time-calibrated tree in BEAST X by applying the same prior on the mtDNA 3^rd^ codon substitution rate derived for *Drosophila melanogaster* (Haag-Liautard et al. 2008). We ran 100,000,000 iterations of Markov Chain Monte Carlo (MCMC) with a 10% burn in and recorded a sample every 10,000 steps. The resulting log file was assessed for convergence in Tracer v1.7.2 before a consensus tree was generated with TreeAnnotator v10.5.0. The phylogenetic tree was visualized in iTOL.

### Characterization of Endosymbiotic *Wolbachia* Strains

We explored both *w*AlbA or *w*AlbB *Wolbachia* infection density in *Ae. albopictus* from Panama in comparison to worldwide populations(Schmidt et al., 2026). To account for variation in sequencing performance, the *Wolbachia* infection density for each sample was normalised by calculating the log ratio of sequencing depth to the depth of the mitochondrial genome. To explore the population structure of *Wolbachia*, we generated a PCA of genotype variation using 115 SNPs segregating in the Panama samples. We used the Smartsnp v1.1.0 R package (Herrando-Pérez et al., 2021) to scale and project our data onto the PCA space of the previously sequenced global samples to account for differences in sequencing depth and genome coverage (Schmidt et al., 2026). PCA plots were generated with the Python package plotly.express (Ploty Technologies Inc, 2015) .

## Results

### Sample sequencing

Mitogenomes from *Ae. albopictus* collected in 2016 and 2017 were successfully sequenced with a high sequencing read number for 57 out of 68 sample libraries (Supplementary Table 1, Figure 1). These had a minimum genome coverage of 96% and a minimum sequencing depth of 830 X. The median sequencing depth was 4517 X. Samples were aligned to the *Ae. albopictus* mitogenome reference (KR068634.1). A total of 64 Single Nucleotide Polymorphisms (SNPs) segregated within the Panama samples. The *Wolbachia* genomes generated for each sample had a median coverage depth of 10 X and a mean genome coverage of 35%.

### Invasion history

To investigate the invasion history of *Ae. albopictus* into Mesoamerica, the mitochondrial genomes from Panama were analysed together with previously generated data from worldwide locations (Battaglia et al., 2022; Schmidt et al., 2026). The number of maternal invasion events that gave rise to our sample of mosquitoes from Panama was estimated by modelling Bayesian phylogeographic diffusion using Panama and the rest of the world as the two traits specified in a CTMC model with a stochastic search variable (Baele et al., 2025; Lemey et al., 2009). This revealed a median 95% HPD count of four to six transitions into Panama from the worldwide distribution of *Ae. albopictus*. This may be an underestimate if lineages found in Panama coalesce in a population elsewhere that may not have been sampled.

Given that *Ae. albopictus* were first documented in Panama 2002, an alternative way to estimate the number of mitochondrial lineages that were introduced and present in our sample is to use sequence divergence to identify separate introductions. Assuming 15 generations per year, we expect ∼225 generations to have passed since the first introduction, so ∼450 generations would separate two alleles that shared a common ancestor in 2002. Given an alignment of 16,660 base pairs and a mutation rate of 6.20 x 10^-8^ per site, we can calculate the genome mutation rate to be 1.03×10^-3^ per generation. From the binomial distribution, we can calculate that there is only a ∼1% chance of three or more mutations over 450 generations. Conservatively ignoring purifying selection, we can therefore conclude that pairs of sequences differing by this number of SNPs likely represent separate introductions. Using UPGMA clustering, we found 12 groups of sequences that differed by three or more SNPs, and providing an approximate lower bound on the number of females introduced into Panama (Supplementary Figure 1). Therefore, two independent methods suggest multiple introductions into Panama, likely involving a small number of individuals.

To determine the origin of the invasion events into Mesoamerica, we reconstructed the ancestral location of mitochondrial lineages using the Bayesian approach implemented in BEAST X (Baele et al., 2025). The tree revealed three major clades reported by(Schmidt et al., 2026), representing invasive populations from Indonesia and the Philippines in Southeast Asia (Clade I), temperate East Asia (Clade II) and tropical South East Asia (Clade III) (Figure 2). Clade II can be further divided into lineages from Japan and China (Clade II-Japan and Clade II-China), both of which contain *Ae. albopictus* from Panama and Europe. In contrast, samples from Mexico and the southern USA are only found within Clade II-Japan. All these regions trade with Panama and are potential invasion sources.

**Figure 2.**
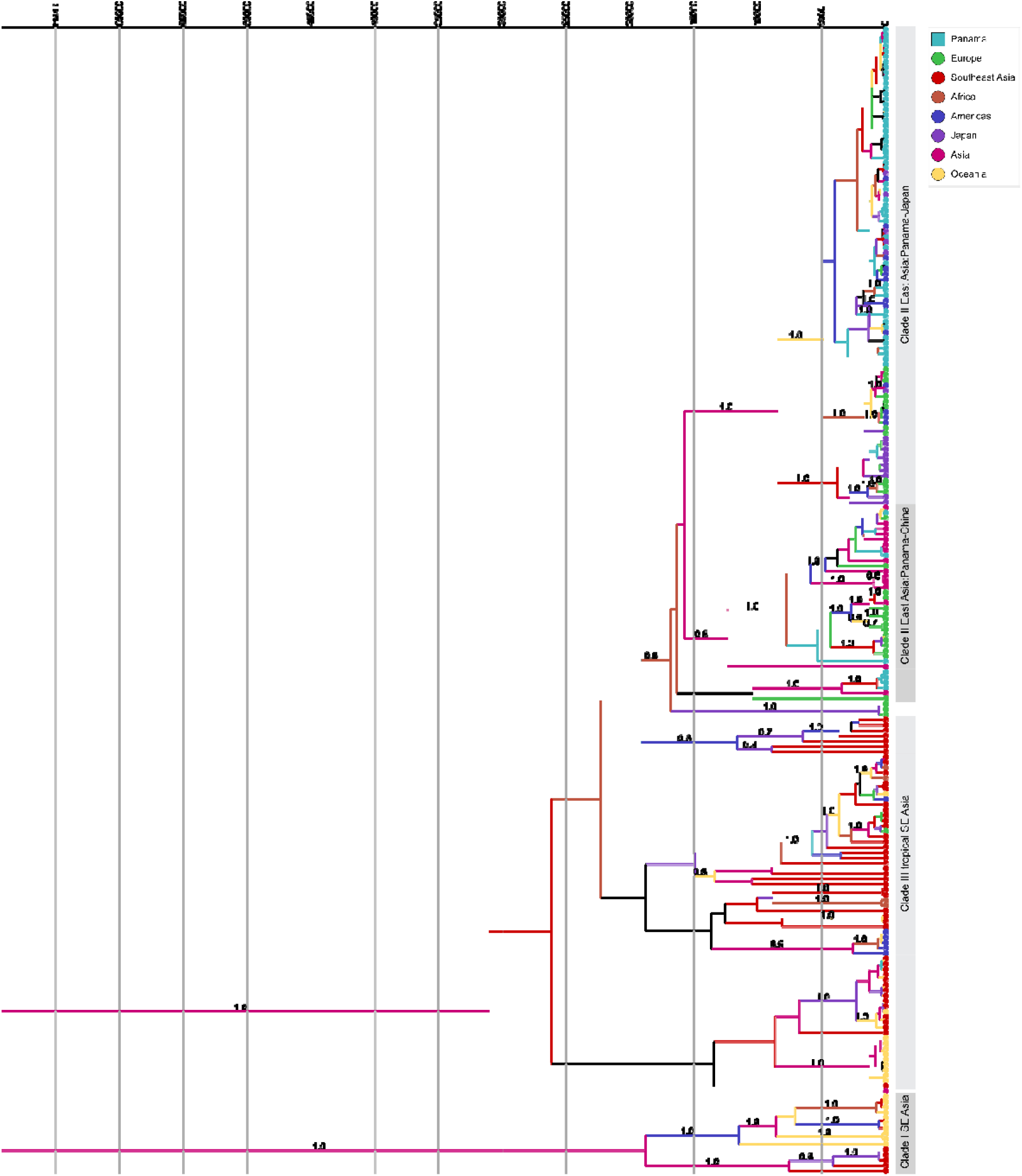
Genealogical tree created in BEAST with samples colored by their location of origin. Branches are coloured by the derived ancestral state location. The placement of divergence time estimates for the Clade II Panama-Japan and Clade II Panama-China based on a 3^rd^ codon position substitution rate are annotated with a black square. Branches with posterior estimates above 0.5 are annotated.

To investigate whether *Ae. albopictus* invaded from Asia directly or indirectly via Europe and/or the USA, we applied a Bayesian phylogeographic diffusion model as previously described with the discrete traits set to the continental origin of each sample. Two separate analysis runs revealed a median of five (95% HPD 3-8) or 6 (95% HPD 4-9) transition counts moving from the Americas to Panama while other regions exhibited a median count of zero, suggesting invasion of populations with Asian ancestry occurred from elsewhere in the Americas. Setting the specific country of origin as the discrete trait yielded inconsistent results, so it is not possible to more precisely identify the source of the invasion.

To estimate the divergence times of the mitogenome lineages introduced into Panama, we used the time calibrated tree described above. We estimated that *Ae. albopictus* from Panama in Clade II-Japan were estimated to have diverged from a common ancestor 4,946 generations ago—allowing for 15 generations per year (Gatt et al., 2009) this places the estimate at 326 years ago. The Panama lineages in Clade II-China diverged from a common ancestor 17,336 generations ago, or 1,144 years ago. As *Ae. albopictus* only began its worldwide expansion from Asia in the 1980’s (Benedict et al., 2007), these estimates reflect the additional time to coalescence of lineages after ‘returning’ to their geographical origin.

### Mitochondrial lineages likely arrived via maritime shipping routes

To explore whether invasion into Panama is likely to have occurred through the shipping canal or overland, we increased our sample size within Panama and added data from Costa Rica, which is the only country with road links to Panama. We generated and aligned 24 mitochondrial *COI* sequences of *Ae. albopictus* collected from Costa Rica in 2019 and 2020 (Supplementary Table 2, Figure 1) and combined these with 117 published Panamanian *COI* sequences from 2014-15 (Eskildsen et al., 2018) and the 59 *COI* sequences from the mitogenomes generated above. In a neighbor-joining tree we observed ten *COI* haplogroups (Figure 3, Supplementary Figure 2), including eight previously described from Panama(Eskildsen et al., 2018). Most sequences from Costa Rica formed a new haplogroup, H9, which was absent from Panama. This had low bootstrap support and was equally genetically similar to haplogroups H3 and H1, both widespread in Panama, Haplogroup H9 had one SNP diagnostic to each of these haplogroups and therefore a pairwise identify to each group of 99.78%. It is known from limited sampling that H3 was historically present in La Virgen in northeast Costa Rica in 2012 (Eskildsen et al., 2018; Futami et al., 2015). However, the H1 haplogroup present in Panama is likely ancestral to both haplogroups H3 and H9, since the two SNPs diagnostic to each H3 and H9 are ancestral (i.e., shared with the *Ae. aegypti* outgroup) in this haplogroup. Haplogroup H3 included 53 out of 59 *COI* sequences extracted from the Panama mitogenomes. This included all individuals in Clade II-Japan and three in Clade II-China. In 2015, haplogroup H3 was restricted to areas close to the major highways of Panama (Eskildsen et al., 2018), suggesting it may have expanded in recent years (Supplementary Figure 3). Haplogroup H8 was found in both countries. However, while it is widespread in Panama (Eskildsen et al., 2018), within Costa Rica it is restricted to areas close to the Panama border, suggesting invasion overland from Panama. Other haplogroups including H1, H2, H4, H5, H6, H7 and H10 are found in Panama but not the Costa Rican sample. Together these results suggest that *Ae. albopictus* was introduced through the shipping canal rather than along highways from Costa Rica.

**Figure 3.**
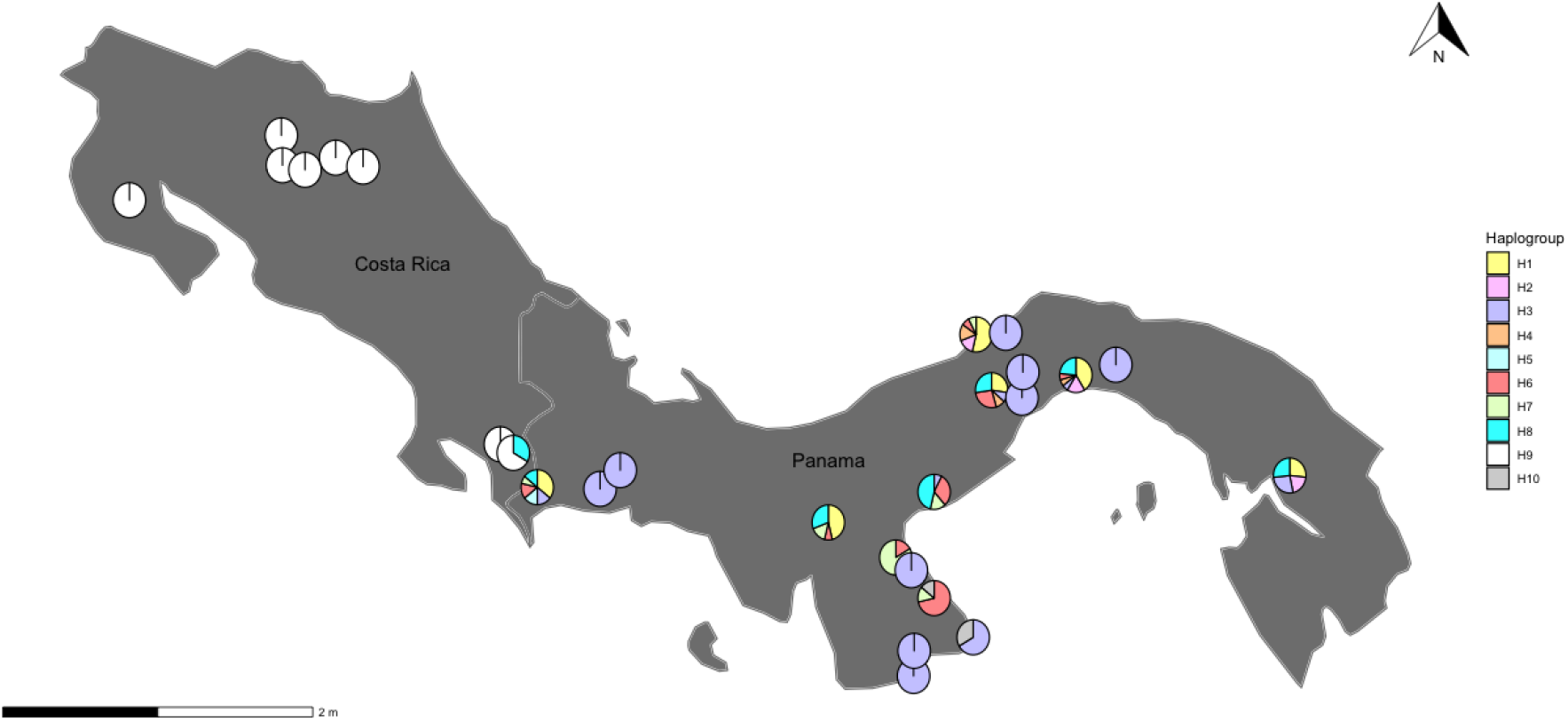
Map of *Ae. albopictus* sampling sites across Panama and Costa Rica for which *COI* sequences were generated. The proportion of individuals with each *COI* haplogroup is displayed.

### *Wolbachia* infection

*Ae. albopictus* is infected with two strains of *Wolbachia*, with the *w*AlbB typically having the highest density within host tissues. To compare the relative density of *Wolbachia* infection across Panama, we calculated the log_2_ ratio of mtDNA read depth to *Wolbachia* read depth and plotted this alongside densities for worldwide locations (Schmidt et al., 2026). The relative density of wAlbB infection was approximately one seventh (median = -8.64 (-10.963 to -2.649)) of the density observed in other worldwide locations (median=-5.855 (-15.458 to -1.1985)) and differed across regional using multiple populations for comparison from Asia (Anova: *F*=28.031, *P*=0.000), South East Asia (Anova: *F*=6.119, *P*=0.003) and Oceania (Anova: *F*=49.702, *P*=0.000) (Figure 4A). The relative densities of *w*AlbA were also markedly much lower (median = -10.66 (-18.062 to -8.576)) at one sixth of the density observed in other worldwide locations (median = -6.052 (-17.826 to -2.723)) (Figure 4B). Anova analysis confirmed that *w*AlbA relative density values differed from those previously reported from multiple countries in Asia (Anova: F= 70.743, P=0.000), South East Asia (Anova: F= 27.738, P=0.000) and Oceania (Anova: F= 48.031, P=0.000).

**Figure 4.**
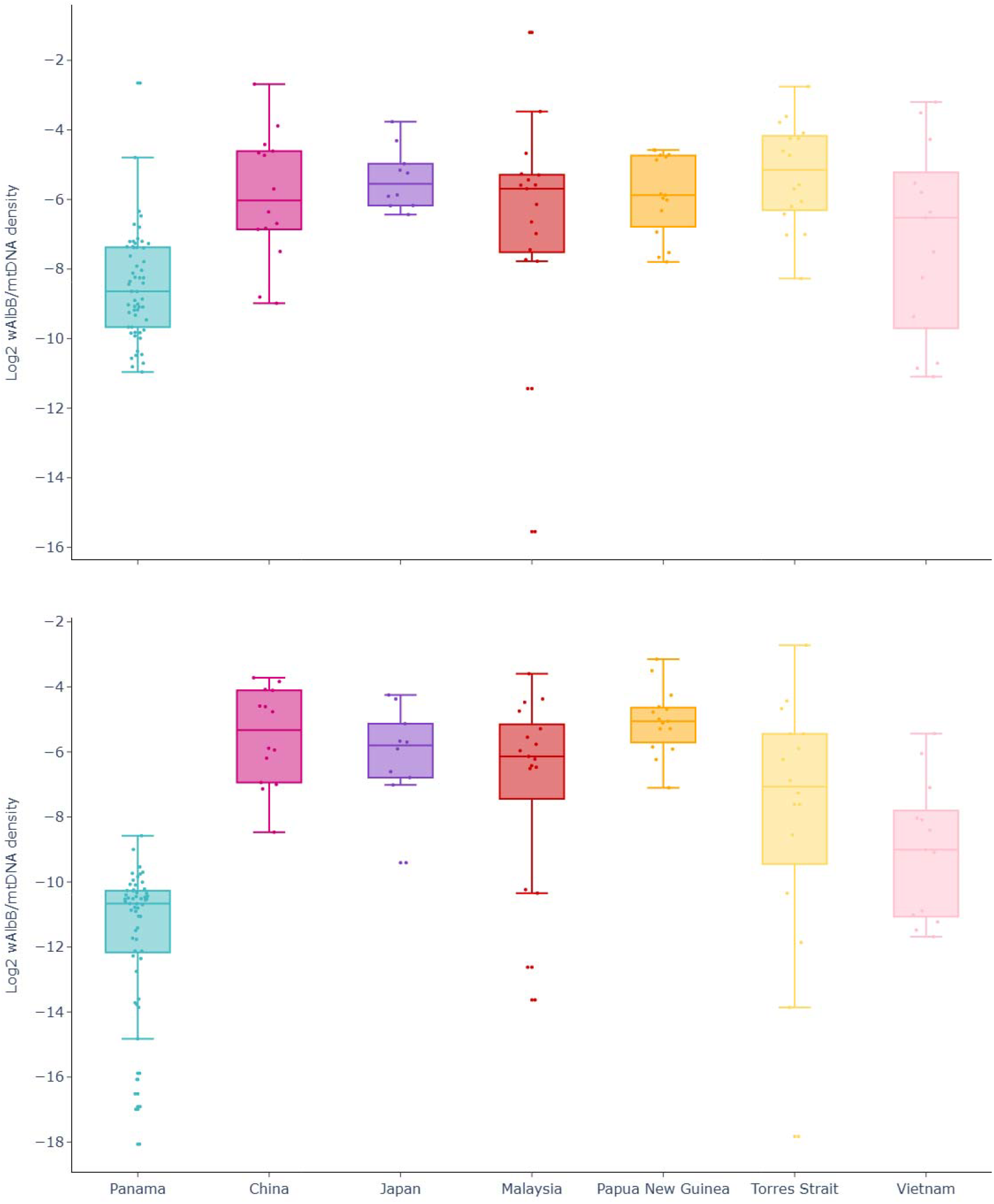
Relative density of (A) *w*AlbB and (B) *w*AlbA relative infection density in *Ae. albopictus* from Panama.

We also compared *Wolbachia* relative density across the different locations sampled in Panama and found that *w*AlbB varied across the country (Anova F= 2.343, P=0.045) whereas *w*AlbA did not (Anova: F= 0.707, P= 0.646). To investigate whether urbanicity impacts the relative density of *w*AlbB, we used a one-way Anova to compare the most urban (Panama City and Colon) versus the most rural (Darien, Chiriqui and Azuero West) locations sampled (Supplementary Figure 4). Densities were significantly higher for some but not all comparisons including Panama City compared to Darien (t=2.637,d.f=4.772,p=0.048) and the city of Colon in comparison to both Chiriqui (t=2.413,d.f=15.739,p=0.028) and the Darien (t=3.943,d.f=6.753,p=0.006).

The population structure of maternally transmitted *Wolbachia* have been found to be congruent with the phylogeographic history of the mosquito (Schmidt et al., 2026). The range of sequencing coverage of the Panama *Wolbachia* genomes was variable (0.13 to 516 X) and differed from previously reported genomes (Schmidt et al., 2026). Therefore, to avoid any bias due to differences in sequencing coverage and depth, we performed a principal components analysis (PCA) where the partial genome sequences of Panama samples were projected onto the PCA space of worldwide *Wolbachia* samples. We removed some samples from the global sample of genomes that strongly influenced the PCA: *w*AlbB from *Ae. albopictus* from Timor and Torres Strait that composed a divergent clade on the phylogenetic tree of worldwide mitogenomes (Clade III) and one sample from China (China145CP031221). Once these divergent samples were removed, *w*AlbB from Panama clustered closely to the Asian samples (Figure 5). The majority of *w*AlbB isolated from Panama, formed a distinct group that was genetically similar to Japan. However, some individuals were more dispersed on the plot, and clustered together with *w*AlbB from China. Findings are therefore generally congruent with findings from the phylogenetic analysis of *Ae. albopictus* mitogenomes and suggest that *w*AlbB genotypes were introduced into Panama from separate invasion routes involving mosquitoes of Japanese and Chinese ancestry.

**Figure 5.**
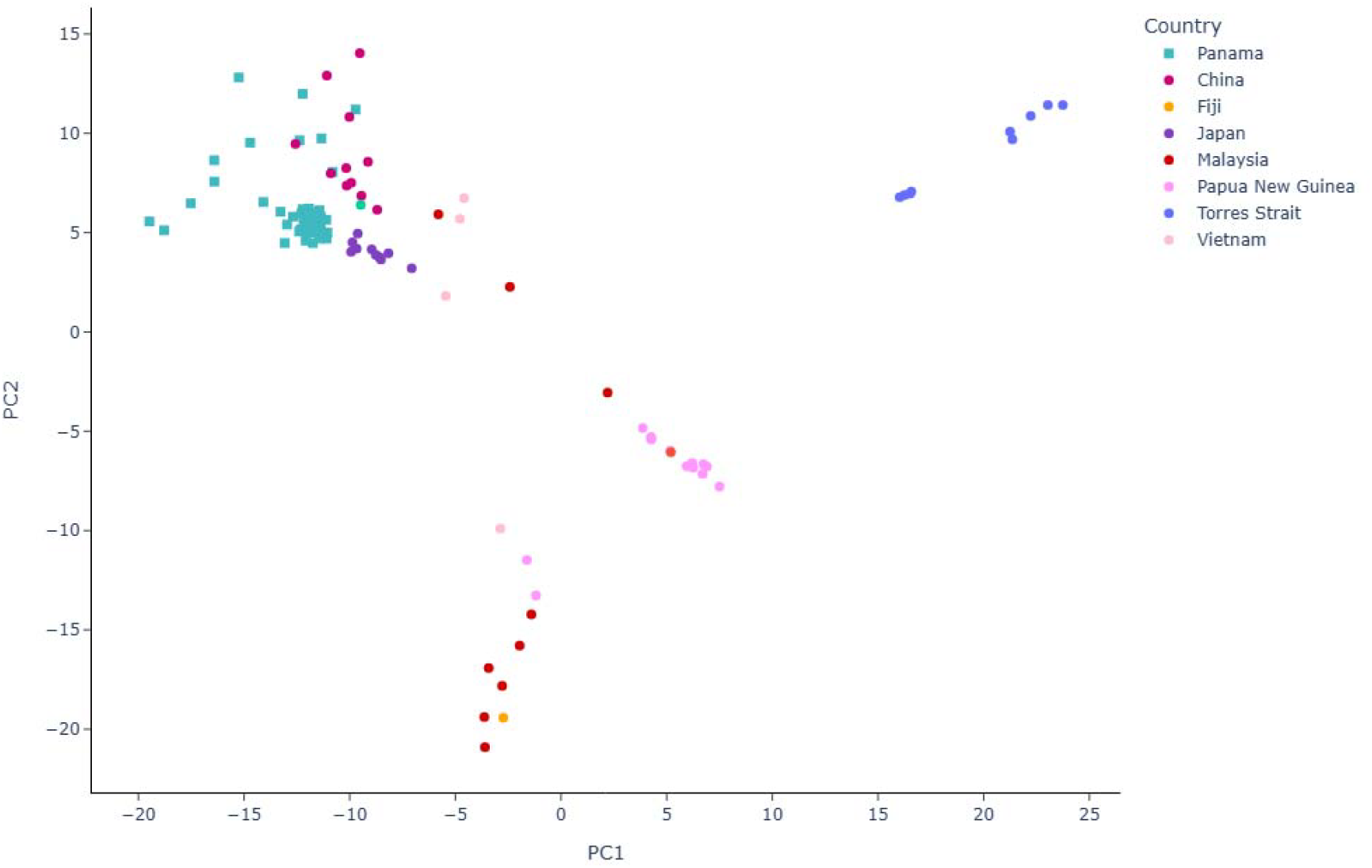
PCA of *w*AlbB genotypes isolated from *Ae. albopictus*. The samples from Panama are projected onto the PCA space of the Asian samples.

## Discussion

By investigating the ancestry of *Ae. albopictus* within Mesoamerica we have uncovered invasion sources and colonization routes. Our findings suggest that maritime invasion via the Panama shipping canal is an important introduction point for *Ae. albopictus* into Mesoamerica. The trade in used tyres is known to facilitate the dispersal of mosquito eggs in a desiccated state, which then hatch on arrival when exposed to rainwater. The Panama Canal has been suggested as the likely source of first invasion into Panama because *Ae. albopictus* was first recorded from the near the canal at Juan Diaz, Panama City in 2002(Miller & Loaiza, 2015).

We found genomic evidence that *Ae. albopictus* have invaded Panama multiple times from populations that traced their ancestry back to Asia, with multiple introductions from two genetically differentiated mitochondrial subclades (Clade II-Japan and Clade II-China). The majority of samples from Panama fell into a subclade containing both samples from the USA and Japan. Our analysis suggests these introductions were not direct from Asia, but instead were first introduced elsewhere in the Americas, and from there transported to Panama. This is likely because the USA is historically the most prolific user of the Panama Canal while Japan’s use is secondary(Reiter, 1998). Furthermore, it is known that *Ae. albopictus* first invaded the southern states of the USA in the late 1980’s and that this was most likely through the importation of tyres and bamboo stumps from Japan(Garcia-Rejon et al., 2021; Hawley et al., 1987). It has also previously been shown that *Ae. albopictus* from the USA share ancestry with Japan based on mitogenomes (Battaglia et al., 2022), supporting our finding that they are genetically similar. We found that one of *Ae. albopictus* from Mexico also fell into the subclade with samples from Japan, USA and Panama. It is unsurprising this sample would be closely related to *Ae. albopictus* from the USA, since it is reported to have invaded Mexico in 1988 close to the USA border(Ortega-Morales et al., 2022). Furthermore, a previous study found both Panama and Mexico to share ancestry with the USA based on microsatellites (Vega-Rúa et al., 2020), suggesting both are derived from the USA. Overall, it is likely that most *Ae. albopictus* in Panama were sourced from the USA and have Japanese ancestry. We found that few individuals from Panama fell into a subclade (Clade II-China) with samples from China and none from the USA. Despite this our ancestral state reconstruction failed to find support for introductions from locations other than the USA.

The evidence suggests that *Ae. albopictus* did not invade Panama overland from Costa Rica. *Ae. albopictus* from Costa Rica and Panama rarely shared *COI* haplogroups in our analysis aside from near the border. The *COI* haplogroup H3 we observed as common across Panama was reported in northeast Costa Rica in 2012(Eskildsen et al., 2018; Futami et al., 2015). However, this is the only record of the haplogroup in Costa Rica, and it was not found in our data. Instead most *COI* haplogroups were unique to Panama, which supports invasion through the Panama Canal rather than from Costa Rica. (Eskildsen et al., 2018)

Most of the mitochondrial lineages in Mesoamerica trace their ancestry to temperate regions in Japan or China. Assuming that these lineages are not found in unsampled tropical regions, this is unexpected given that mosquitoes from tropical regions tend to have tropical origins (Paupy et al., 2009). Furthermore, mosquitoes from temperate regions are expected to be sub optimally adapted to tropical climates. Adaptation to the tropics may be further hampered in introduced locations due to lower genetic diversity (Estoup et al., 2016). However, frequent migration can boost genetic diversity and evidence suggests that *Ae. albopictus* can rapidly adapt to different environments(Medley et al., 2019). For example, the ability of *Ae. Albopictus* to undergo diapause in response to cold climate conditions is a strong characteristic of temperature regions while lacking or reduced from tropical populations(Lounibos et al., 2003). However, *Ae. albopictus* originating from temperate Japan in the subtropical southern states of the USA have reduced their diapause response on invasion(Lounibos et al., 2003; Medley et al., 2019). This adaptation is likely due to the fitness cost imposed by diapause which can impact egg viability(Sherpa et al., 2022). If invasions into Mesoamerica are sourced from subtropical populations unsampled from within the USA, it is possible these are better adapted to tropical climates than expected by populations invading directly from a temperate region such as Japan.

Invasion events have the potential to introduce novel strains of *Wolbachia* that impact population control. We observed that mosquitoes have similar *Wolbachia* to China and Japan, supporting the observation of *Ae. albopictus* mtDNA in Panama with Chinese and Japanese ancestry. That the phylogeographic patterns of mtDNA and *Wolbachia* are closely matched is not surprising given that they are similarly inherited through the maternal line. Furthermore, it has previously been shown that the phylogenies of mosquito mitogenomes and *Wolbachia* genomes were congruent (Schmidt et al., 2026). A deep understanding of the geographic distribution of naturally circulating *Wolbachia* strains is critical to successful implementation of *Wolbachia* population suppression strategies. This is because successful population control with *Wolbachia* relies on the interaction between mosquito fitness, cytoplasmic incompatibility and its ability to reduce disease transmission(Montenegro et al., 2024). The presence of or introduction of a naturally occurring *Wolbachia* strain into a population has the potential to compromise release strategies if the strain has a fitness benefit or pattern of cytoplasmic incompatibility that promotes its spread. For example, the laboratory modified *Wolbachia* strain *w*Mel has bidirectional cytoplasmic incompatibility with the naturally occurring strains *w*AlbA and *w*Alb and compromises the efficacy of releases(Hoffmann et al., 2015). This has led to the development of triple superinfected mosquitoes which can overcome the barriers imposed by local *Wolbachia* strains (Zheng, Zhang, Li, Yang, Wu, Liang, Liang, Pan, Hu, & Sun, 2019). Given the genetic similarity of recently invaded *Ae. albopictus* and its associated *Wolbachia* to source populations, it is possible that biological knowledge aiding population control could be transferred across geographical boundaries. For example, successful *Wolbachia* control strategies applied to source populations, may be similarly effective in places where the mosquito has only recently invaded.

We found that the relative density of *w*AlbB infection in *Ae. albopictus* from Panama was higher than *w*AlbA, consistent with findings from elsewhere suggesting lower densities are found due to differences in tissue tropism and replication rates (Chuchuy et al., 2025; Dutton & Sinkins, 2004). The relative density of both *w*AlbB and *w*AlbA was lower in Panama compared to other worldwide locations. Variation in *Wolbachia* density between countries has been reported before(Schmidt et al., 2026). It is known that the host genotype (Padde et al., 2023) and environmental factors like diet and temperature influence the *Wolbachia* densities (Mouton et al., 2007). Temperatures are often at the top end of the range *Ae. albopictus* can tolerate in Panama, and high temperatures are associated with lower densities of *Wolbachia* infection. However, *w*AlbB is less impacted by high temperature than some other *Wolbachia* strains (Mouton et al., 2006; Ross et al., 2017, 2019). Diet is also important, likely because *Wolbachia* relies on the nutrients required by the host as the bacteria do not break down amino acids and lipids (Caragata et al., 2014; Dutton & Sinkins, 2004; Padde et al., 2023). If *Ae. albopictus* are encountering nutrient poor environments in Panama, this could reduce *Wolbachia* titer. (Bennett et al., 2021b; Deerman & Yee, 2023; Murrell et al., 2011). Regardless of the causes, *Wolbachia* infection densities impact on vertical transmission and cytoplasmic incompatibility, which may be reduced or even halted at low densities (Calvitti et al., 2015; Ross et al., 2019). Furthermore, the ability of *Wolbachia* to block viral replication in the host is compromised when densities are low (Ant et al., 2018; Lu et al., 2012; Padde et al., 2023). These findings highlight the need to further characterize the influence of climatic and environmental parameters on *Wolbachia* density and to assess how these impact the successful spread of *Wolbachia* and reduction of virus transmission in both the laboratory and field.

Overall, we have found that the Panama Canal is an important introduction point of invasive *Ae. albopictus*. Given the ecological similarity of other invasive *Aedes* vectors, findings suggest it may also be an important surveillance site for the early detection of species currently expanding across the Americas but not yet present in the Isthmus, such as *Aedes vittatus* (Ng’eno et al., 2025). That we observed multiple introductions of *Ae. albopictus* into Mesoamerica is concerning because migration can increase genetic diversity, boost population sizes, introduce adaptations that impact on public health and introduce new viral strains(Lindh et al., 2019; Medley et al., 2019). For example, E1 and E2 glycoprotein mutations identified in *Ae. albopictus* increases mosquito infectivity and its ability to transmit Chikungunya virus with the potential for geographical spread (Tsetsarkin et al., 2009). Furthermore, *kdr* substitutions at the voltage gated sodium channel which confer broad range insecticide resistance are yet to be observed in *Ae. albopictus* from Panama(García et al., 2024). However, this adaptation is likely to rapidly spread on introduction to compromise pyrethroid based spraying campaigns as occurred for *Ae. aegypti* (Tuñon et al., 2024). Therefore, the arrival of adaptive alleles of public health concern could be monitored at invasion points through genomic surveillance. Timely detection of this type of adaptive threat followed by proactive intervention has the power to generate a positive outcome for disease management.

## Conclusion

Our study reveals that *Ae. albopictus* populations in Panama arise from multiple recent introductions, predominantly mediated through maritime trade linked to the Panama Canal rather than overland dispersal from neighbouring regions. Integrating complete mitochondrial genomes with maternally co-inherited *Wolbachia* markers provides convergent evidence for distinct invasion pathways tracing back to Asian-derived lineages via the Americas. Notably, the reduced *Wolbachia* densities observed in Panama raise important questions about environmental influences on symbiont dynamics and their downstream effects on arbovirus transmission and control strategies. Together, these findings identify the Panama Canal as a key route for vector introductions into Mesoamerica.

## Acknowledgements

We are grateful to the Panamanian Ministry of Environment (MiAmbiente) for supporting the scientific collection of mosquitoes, and to the Ministry of Health (MINSA) for providing historical data on Aedes distribution from 2005 to 2017 across the country. We also thank the Smithsonian Tropical Research Institute (STRI) and the Instituto de Investigaciones Científicas y Servicios de Alta Tecnología (INDICASAT) for their administrative support, technical guidance, and logistical management. Finally, we thank the anonymous reviewers for their valuable comments and suggestions. We thank all members of the Costa Rican National Vector Control Program who kindly helped gathering the samples for this study.

## Data Accessibility Statement

The sequencing data will be submitted by time of publication. All generated sequences were deposited in the NCBI Sequence Read Archive under accession numbers XXXXXX–XXXXXX and GenBank under accession numbers XXXXXX– XXXXXX.

## Benefit-Sharing Statement

A research collaboration was developed with scientists from the countries providing samples for sequencing, all collaborators are included as co-authors and the results of research have been shared. The research concerns the study of an arboviral vector of public health concern.

## Author Contributions

Conceptualization: KLB, WOM, LFC, FJ, and JRL. Investigation (fieldwork): JGA, GD, RMR, and LFC (Costa Rica); KLB, JIL, and JRL (Panama). Investigation (laboratory and taxonomy): LFC, JIL, and JRL. Data curation: KLB, TS, JPD, JGA, GD, RMR, LFC, FJ, and JRL. Visualization: KLB, TS, JPD, LFC, FJ, and JRL. Formal analysis: KLB, TS, JPD, LFC, FJ, and JRL. Writing – original draft: KLB. Writing – review & editing: TS, JPD, JGA, GD, RMR, LFC, JIL, WOM, FJ, and JRL. All authors have read and agreed to the published version of the manuscript.

## Ethical approval

Mosquito collection in Panama was authorized by the Ministry of Environment (MiAmbiente) under permit number ID 8-447-900-PAN.This study was carried out in accordance with Article 7 from Law 9234 for biomedical research which grants the Epidemic Surveillance Division (Vigilancia de la Salud) of Costa Rica□s Ministry of Health (Ministerio de Salud) the ability to perform activities with the dual goal of surveillance and research, which are exempted from the approval of an Institutional Research Board, as these efforts are deemed essential for health policy planning and decision making, and do not release individually identifiable data.

