## Supplementary Material for "Invasion history of *Aedes* (Stegomyia) *albopictus* into Mesoamerica based on mitogenomes and *Wolbachia* symbionts: Multiple introductions with temperate origins"

**Supplementary Tables**

Supplementary Table 1. The number of mitogenomes sequenced from *Ae. albopictus* collected across Panama.

| **Province** | **Location** | **Latitude** | **Longitude** | **Year** | **No. samples** |
| --- | --- | --- | --- | --- | --- |
| Chiriquí | Gualaca | 8.53 | -82.30 | 2016 | 3 |
|  |  |  |  | 2017 | 4 |
|  | David | 8.42 | -82.43 | 2017 | 8 |
| Colón | Sabanitas | 9.35 | -79.80 | 2017 | 6 |
|  | Gamboa |  |  | 2017 | 5 |
| Darién | Metetí | 8.50 | -77.98 | 2017 | 3 |
| Los Santos | La Villa | 7.94 | -80.41 | 2017 | 2 |
|  | Tonosí | 7.41 | -80.44 | 2017 | 5 |
|  | El Cacao | 7.45 | -80.40 | 2017 | 5 |
|  | Pedasí | 7.53 | -80.02 | 2017 | 6 |
| Panamá Oeste | Princesa Mía | 8.97 | -79.70 | 2017 | 5 |
| Panamá | Chepo | 9.16 | -79.10 | 2017 | 5 |

Supplementary Table 2. The number of *Ae. albopictus* from Costa Rica with newly generated cytochrome oxidase I (COI) sequences for analysis.

| **Province** | **Location** | **Latitude** | **Longitude** | **Year** | **No. samples** |
| --- | --- | --- | --- | --- | --- |
| Punta Arenas | Coto | 8.64 | -83.01 | 2019 | 6 |
|  | Rio Claro | 8.69 | -83.07 | 2019 | 1 |
| Guanacaste | Nicoya | 10.14 | -85.46 | 2019 | 2 |
| Heredia | La Virgen | 10.40 | -84.13 | 2020 | 2 |
|  | Sarapiquí | 10.45 | -84.03 | 2019 | 2 |
|  | Yacare | 10.45 | -84.03 | 2019 | 1 |
|  | Horquetas La Victoria | 10.33 | -83.95 | 2019 | 1 |
|  |  |  |  | 2020 | 2 |
| Alajuela | Platanar | 10.42 | -84.47 | 2019 | 1 |
|  | Pital | 10.45 | -84.32 | 2019 | 1 |
|  | Florencia | 10.36 | -84.48 | 2019 | 1 |
|  | Alajuela | 10.54 | -84.48 | 2019 | 3 |
|  | Aquas zarcas | 10.38 | -84.33 | 2019 | 1 |

**Supplementary Figures**

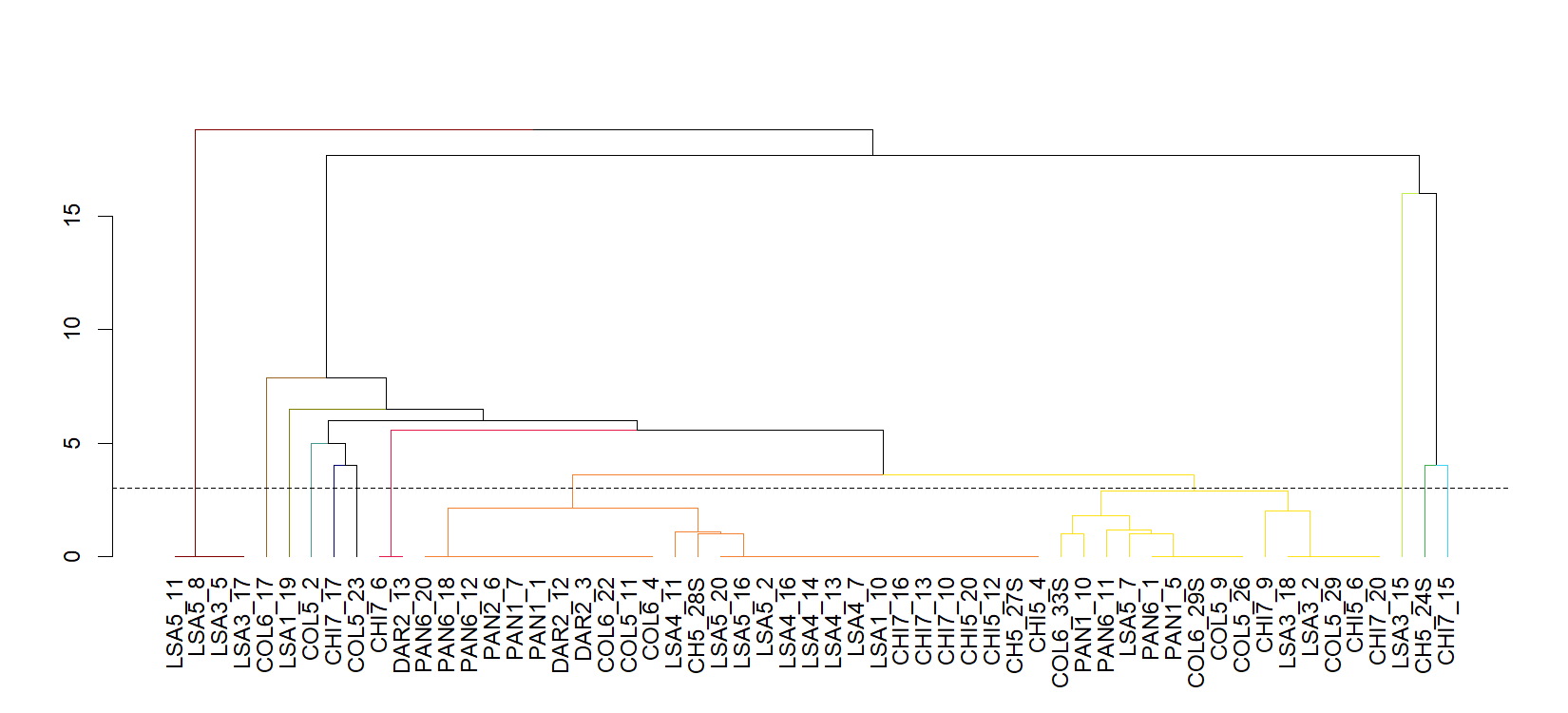

Supplementary Figure 1. A dendrogram assigning individuals to 12 UPMGA groups based on a minimum of two SNP differences as indicated by the dotted line. Dendrogram branches are coloured by UPMGA group.

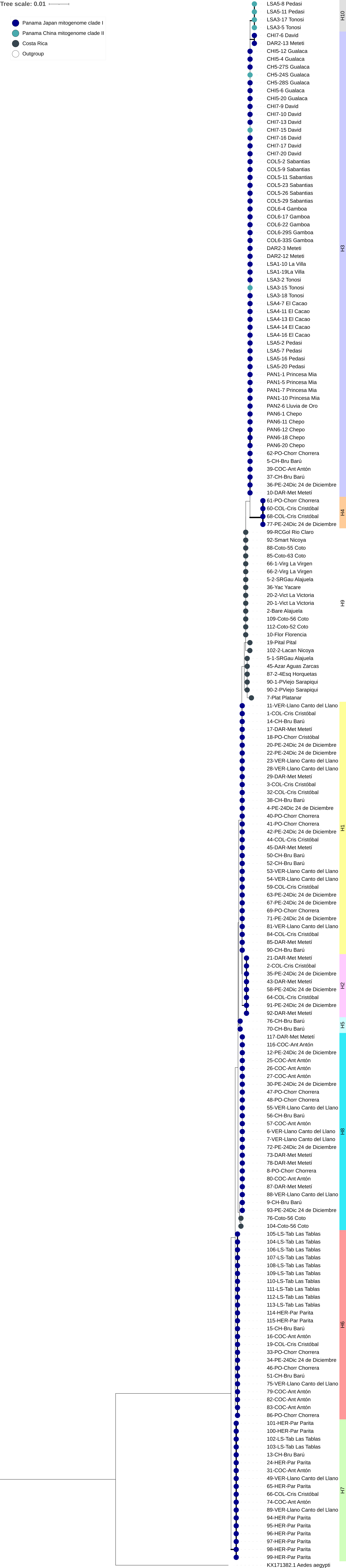

Supplementary Figure 2. Neighbor-joining tree of *COI* sequences with haplogroups annotated.

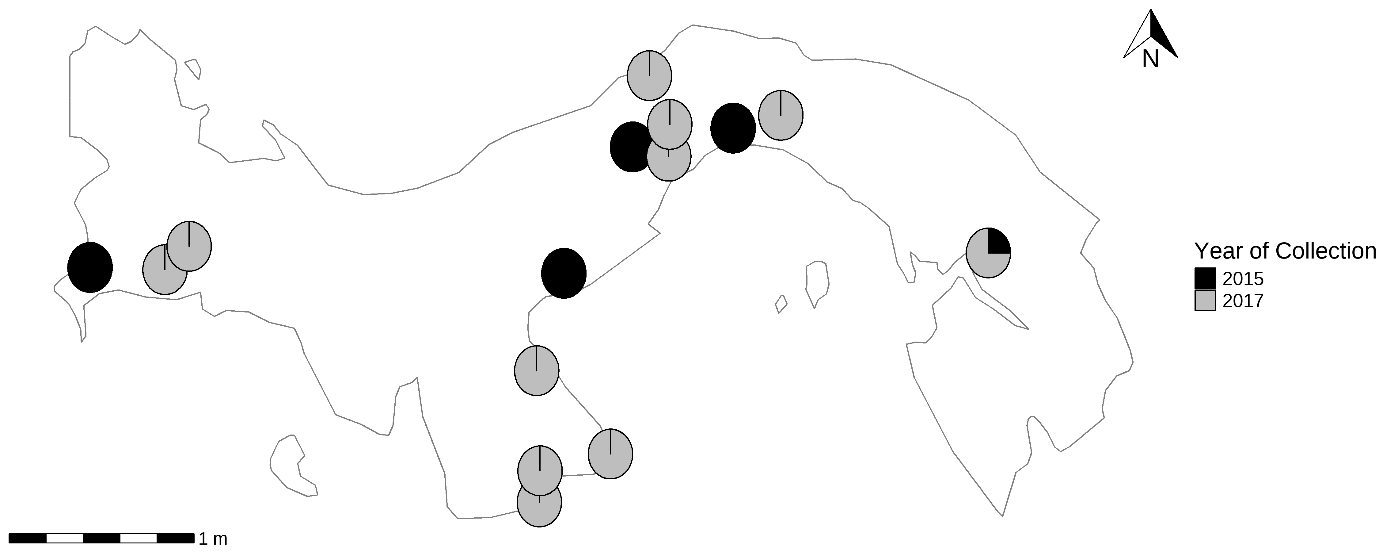

Supplementary Figure 3. Map showing the distribution of haplogroup H3 across Panama in 2015 and 2017.

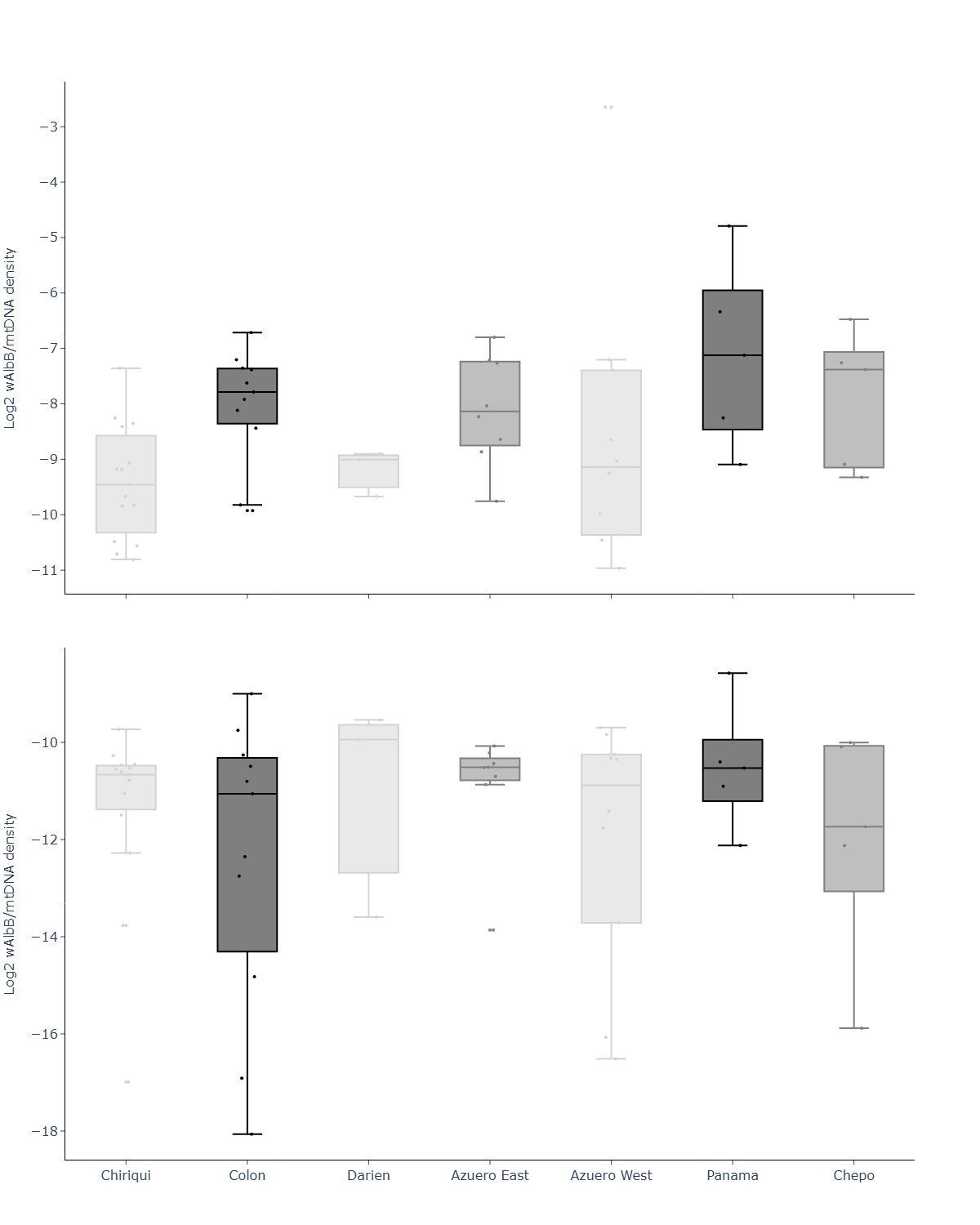

Supplementary Figure 4. Relative density of (a) *w*AlbB and (b) *w*AlbA relative infection density in *Ae.* *albopictus* from locations in Panama.
